# ARIBA: rapid antimicrobial resistance genotyping directly from sequencing reads

**DOI:** 10.1101/118000

**Authors:** Martin Hunt, Alison E. Mather, Leonor Sánchez-Busó, Andrew J. Page, Julian Parkhill, Jacqueline A. Keane, Simon R. Harris

## Abstract

Antimicrobial resistance (AMR) is one of the major threats to human and animal health worldwide, yet few high-throughput tools exist to analyse and predict the resistance of a bacterial isolate from sequencing data. Here we present a new tool, ARIBA, that identifies AMR-associated genes and single nucleotide polymorphisms directly from short reads, and generates detailed and customisable output. The accuracy and advantages of ARIBA over other tools are demonstrated on three datasets from Gram-positive and Gram-negative bacteria, with ARIBA outperforming existing methods. ARIBA is available at https://github.com/sanger-pathogens/ariba.

## Background

Antimicrobial resistant infections have become one of the leading threats to human health, with a conservative estimate of 700,000 directly attributed deaths per year worldwide [1]. If we do not address this threat, this figure is estimated to rise to 10 million by 2050 [1]. An important component of any strategy to tackle antimicrobial resistance (AMR) is having rapid and accurate methods for identifying markers of resistance. Our understanding of the mechanisms and diversity of AMR is improving, in part due to the increased availability of genome sequence data, with the use of genome sequencing in personalised medicine set to become one key tool in the fight against AMR. However, there are currently few bioinformatics tools that can identify AMR determinants directly from the data produced by widely-used sequencing technologies. The methods that are available are limited in the types of AMR mechanisms they can detect and/or are not scalable to high-throughput environments.

Limitations of existing tools include being available only via web services that are not high throughput, being restricted to a specific set of reference sequences which may not exhaustively represent current knowledge of AMR for all microbial species, requiring assembled genome sequences as input, an inability to identify and interpret singlenucleotide-polymorphism (SNP)-based AMR determinants and having high computational resource requirements. Most tools fall into one of two categories: those that align sequencing reads to a set of reference genes, and those that search for reference gene matches in *de novo* assembled sequences. The widely-used SRST2 [2]is an example of a method based on aligning reads to a set of reference sequences in order to predict the presence of those genes in a sample. KmerResistance [3]employs a similar approach, but uses *k*-mer matching between sequencing reads and reference genes to identify gene presence. Although SRST2 and KmerResistance can be used with custom reference gene sets, they cannot directly identify or interpret variants, such as SNPs that confer resistance, and so are limited to identifying resistance that is conferred by the presence of a gene, or a particular pre-defined allele of a gene. Mykrobe predictor [4]is an extremely fast tool that matches *k*-mers in reads to a reference graph, and although it can identify variants, it is currently limited to *Staphylococcus aureus* and *Mycobacterium tuberculosis*, and it is not possible for users to provide their own databases of AMR determinants with which to interrogate their data.

The majority of other AMR detection tools require assembled sequences as input, which are computationally expensive to generate from reads, and assembly errors or failures caused by the complexity of assembling complete genomes *de novo* can lead to AMR determinants being missed. For these reasons, alignment based approaches have previously been shown to be superior to the use of *de novo* assembled sequences [2,3]for AMR gene detection. Tools that use assembled sequences as input include ResFinder [5], ARG-ANNOT [6], SSTAR [7], and RAST [8]. These methods match assembled sequences to reference genes, usually using the BLAST [9]algorithm, in order to identify AMR genes.

Here we present a new tool, called ARIBA (Antimicrobial Resistance Identification By Assembly), that uses a combined mapping/alignment and targeted local assembly approach to identify AMR genes and variants efficiently and accurately from Illumina paired sequencing reads. Using targeted local assembly considerably reduces the complexity of the assembly process, while providing contiguous gene or nucleotide sequences without the ambiguity of the interpretation of aligned data. ARIBA can easily be provided with custom reference sequence-sets, and includes support for a number of public databases: ARG-ANNOT [6], CARD [10], MEGARes [11], and ResFinder [5]. It distinguishes between sequences that are coding or non-coding, and provides details on each sequence present in the sample. It verifies whether or not identified genes are complete, truncated or fragmented in the sample, and reports SNPs and indels within sequences with interpretations of their effect, such as frameshifts, non-synonymous changes or nonsense mutations. To facilitate easier interpretations of results, ARIBA includes functions to summarise results for multiple samples. These summaries are compatible with the Phandango interactive visualisation tool [12]. If minimum inhibitory concentration (MIC) data are available for samples, ARIBA allows statistical analysis and plotting of MIC against genotype. Beyond AMR, ARIBA can be used more generally to find any input sequences of interest. It provides inbuilt support for the PlasmidFinder [13]and VFDB [14]databases, and functionality for multi-locus sequence typing (MLST) using data from PubMLST [15].

## Results

We developed ARIBA to identify AMR determinants from public or custom databases using Illumina paired read data as input. Figure 1 provides an overview of the approach - complete details can be found in the Methods section and Supplementary Material (Supplementary Figures S1-4). Briefly, reference sequences in the AMR database are clustered by similarity using CD-HIT [16]. Reads are mapped to the reference sequences using minimap [17]to produce a set of reads for each cluster. These reads map to at least one of the sequences in that cluster. The reads for each cluster and their sequence pairs are assembled independently using fermi-lite [18]under a variety of parameter combinations, and the closest reference sequence to the resulting contigs is identified with the program `nucmer` from the MUMmer package [19]. The assembly is compared to the reference sequence to identify completeness and any variants between the sequences using the `nucmer` and `show-snps` programs from MUMmer. The reads for the cluster are mapped to the assembly with Bowtie2 [20] and variants are called with SAMtools [21]. Finally, a detailed report is made of all the sequences identified in the sample, including, but not limited to, the presence or absence of variants pre-defined to be of importance to AMR.

**Figure 1.**
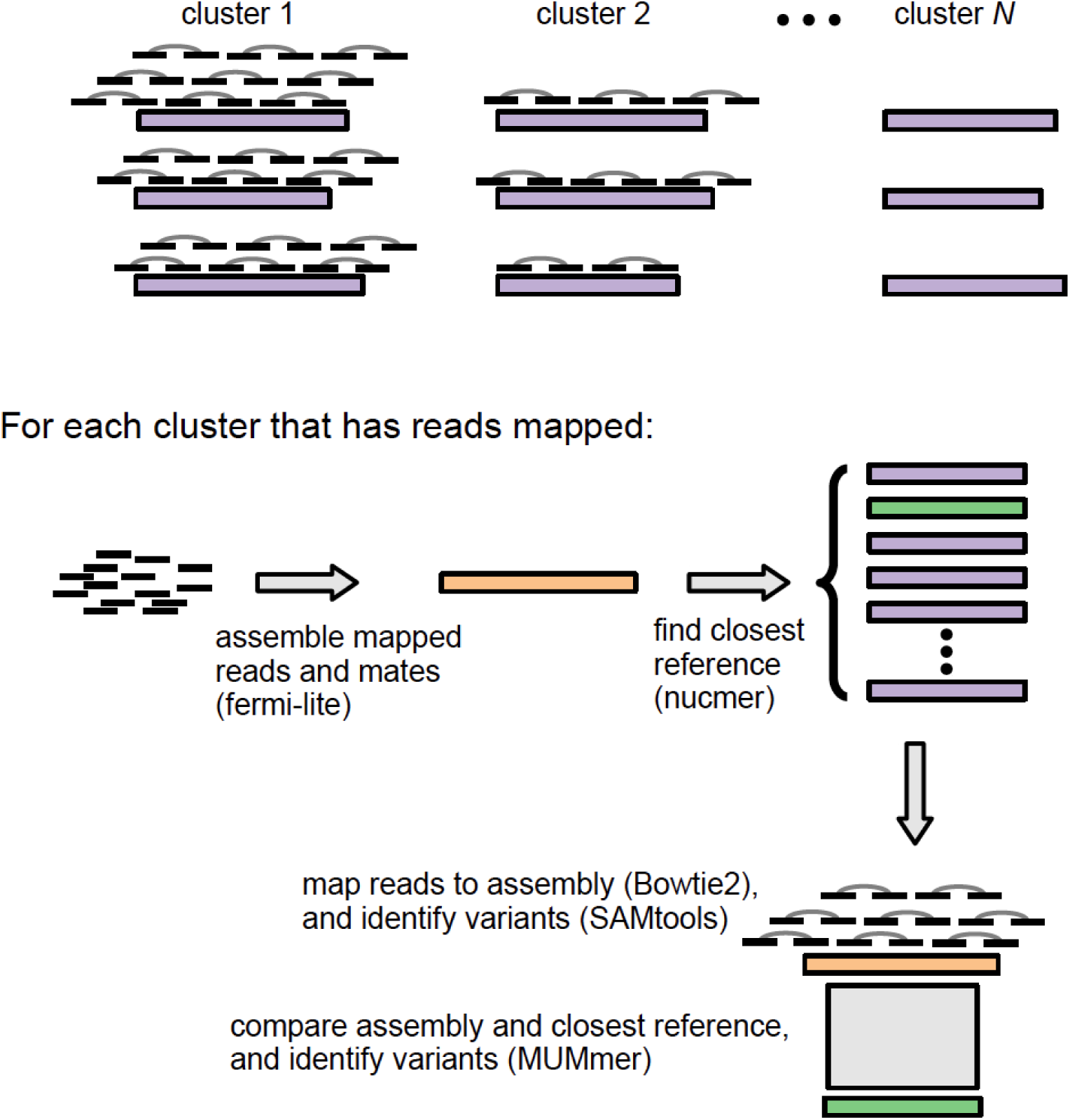
Overview of the ARIBA mapping and targeted assembly pipeline.

The performance of ARIBA was evaluated on three datasets, to illustrate all aspects of its functionality and to benchmark against other available methods. For these comparisons, we focussed on command line tools that can use custom reference data.

### Enterococcus faecium

The first dataset comprises 41 isolates of the Gram-positive bacterium *Enterococcus faecium*, for which the phenotypic resistance to vancomycin is known for each sample [22] (Supplementary Table S1). This dataset, which was used to evaluate SRST2 in its initial publication [23], allowed validation of the accuracy of ARIBA when identifying the presence or absence of genes of interest in each sample. Seventeen of the samples are known to have VanB-mediated resistance to vancomycin (i.e., are vancomycin resistant enterococci, VRE) and the remaining 24 samples are known to be susceptible (i.e., are vancomycin susceptible enterococci, VSE). The phenotypic resistance is due to the presence of an operon comprising up to seven genes van*B*, van*H*, van*R*, van*S*, van*W*, van*X*, and van*Y* [24]. However, *vanW* and *vanY* are not required for resistance [24][25]. We have also used these data to test the sensitivity of ARIBA and other methods at varying depths of read coverage.

First, we used ARIBA and SRST2 to identify the sequence type of each sample, using the *E. faecium* MLST scheme [26]downloaded from PubMLST. Given that MLST loci are chosen to be conserved, single-copy housekeeping genes, identification of MLST should be a simple test for any gene-detection method. As expected, we found that the results generated by both tools were in complete agreement with the known sequence types provided in [22] (Supplementary Table S1). However, the running time of ARIBA was approximately one-fifth that of SRST2 (Supplementary Table S2). Next, ARIBA, KmerResistance and SRST2 were evaluated on all 41 samples using the antimicrobial resistance reference set of genes from SRST2, which is based on ARG-ANNOT and includes all seven genes of interest. All three tools made identical calls on the 17 VRE samples in the *vanB, vanH, vanR, vanS*, and *vanX* genes, except for the choice of closest reference sequence in sample SRR980582, which differed for the *vanB* gene (Supplementary Table S1). Several of the VSE samples contain low-level contamination with *VanA-B* sequences [2]; here, in most cases only ARIBA flagged the genes as partially present at a low read depth, and SRST2 and KmerResistance did not make any prediction about the presence of these genes (Supplementary Table 1).

The remaining differences between the tools were in the identification of *vanW* and *vanY* in the VRE samples. The discrepancies demonstrate the benefits of the detailed output of ARIBA, when compared to the other tools. A complete description of the differences between the output of the three tools is given in the Supplementary Material. For example, in sample SRR980557, SRST2 reported that the *vanW* gene was present but with one SNP (“1snp” in the output), and KmerResistance also reported the gene as present. ARIBA reported a SNP, but provided the further information that it was a nonsense mutation and therefore the gene is likely to be non-functional in that sample.

The effect of read depth was assessed on the 17 VRE samples by uniformly sampling from the reads at depths ranging from 1 to 100X coverage of the vancomycin resistance operon. The total number of calls for the five required resistance genes made by each tool across all 17 samples is shown in Figure 2, and a per-gene breakdown is given in Supplementary Figure S5. KmerResistance appears to be optimised for coverage below 5X, and its ability to call the presence of genes decreases in the range 2X-18X before recovering in the range 50-75X. The ability of ARIBA and SRST2 to identify genes improves with read depth, with ARIBA marginally outperforming SRST2. When partial matches to genes are included, ARIBA and SRST2 become more sensitive at lower coverages and ARIBA becomes more sensitive than KmerResistance (Supplementary Figure S6).

**Figure 2.**
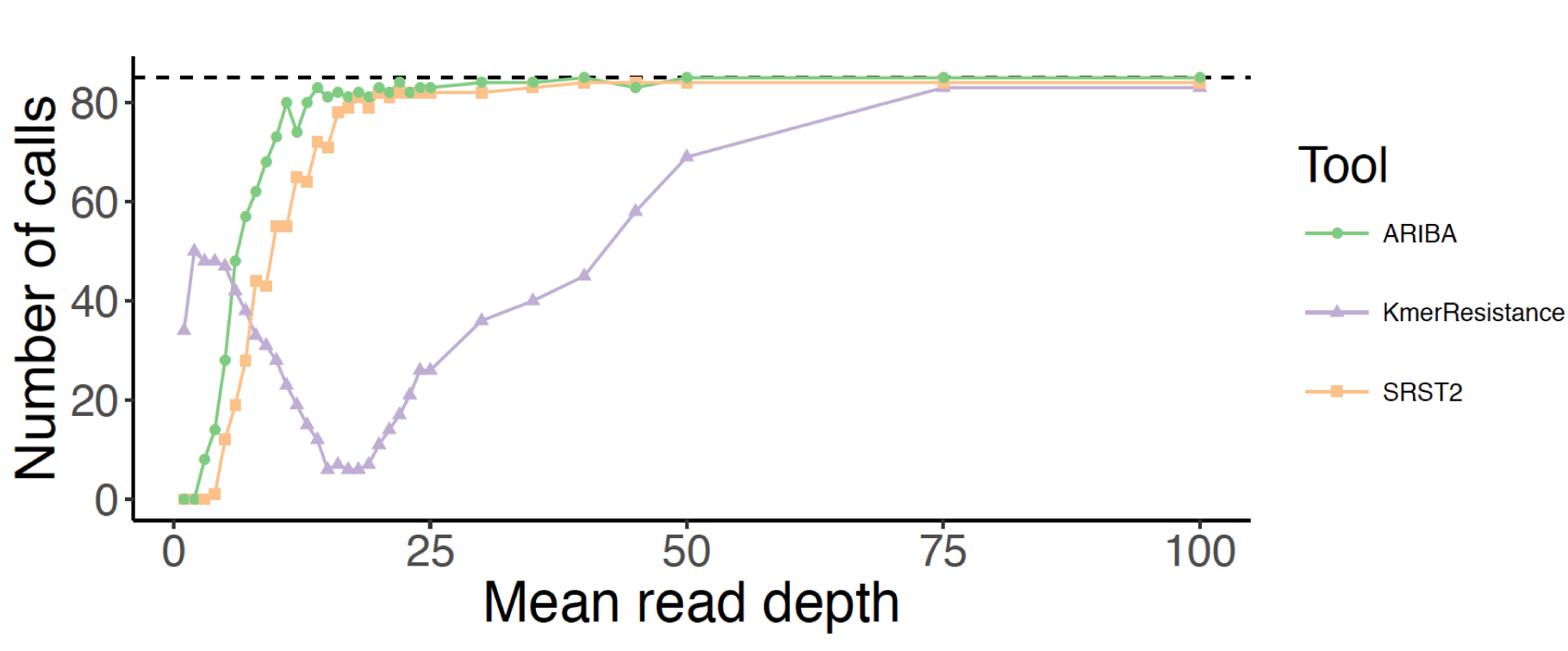
Effect of read depth on the number of gene calls for all five *van* genes in the 17 vancomycin resistant *E. faecium* samples.

### Shigella sonnei

The second dataset, published by Holt *et al.* [23], consists of 130 globally distributed genomes of S. *sonnei* (Supplementary Table S3), a Gram-negative bacterium that is a causative agent of dysentery. The phenotypic resistance profile for a number of antimicrobials is known for each isolate, and is attributable to both acquired resistance genes and SNPs. This enabled a comparison of ARIBA, SRST2, and KmerResistance with the manual method employed in [23], confirming the accuracy of ARIBA for identifying known resistance SNPs as well as the presence or absence of genes of interest. The three tools were run on all 130 samples using the reference database from CARD [10], version 1.1.2. To ensure our results were comparable with those originally reported in Supplementary Table 1 of [23], we manually added those AMR genes listed on page 4 of their supplementary text not already included in the database (Supplementary Table S4). The AMR determinants originally reported in [23]were identified from mapping data, and reported as the proportion of bases in the gene sequence that were covered by reads from each isolate. From these originally reported data, we used a cut-off of ≥ 90% to indicate that a gene was present by their method.

With seven antimicrobials and 130 isolates, there was a potential for 910 AMR calls (identification of a gene, set of genes, or SNP). In 546 cases, no calls were made by any method. 364 AMR calls were made by at least one of the four methods; 60% (218/364) were found by all four methods (Figure 3 and Supplementary Table S5). Overall, this results in an agreement between the four methods of 84%: (218 calls in agreement + 546 non-calls in agreement) / (910 potential AMR calls).

**Figure 3.**
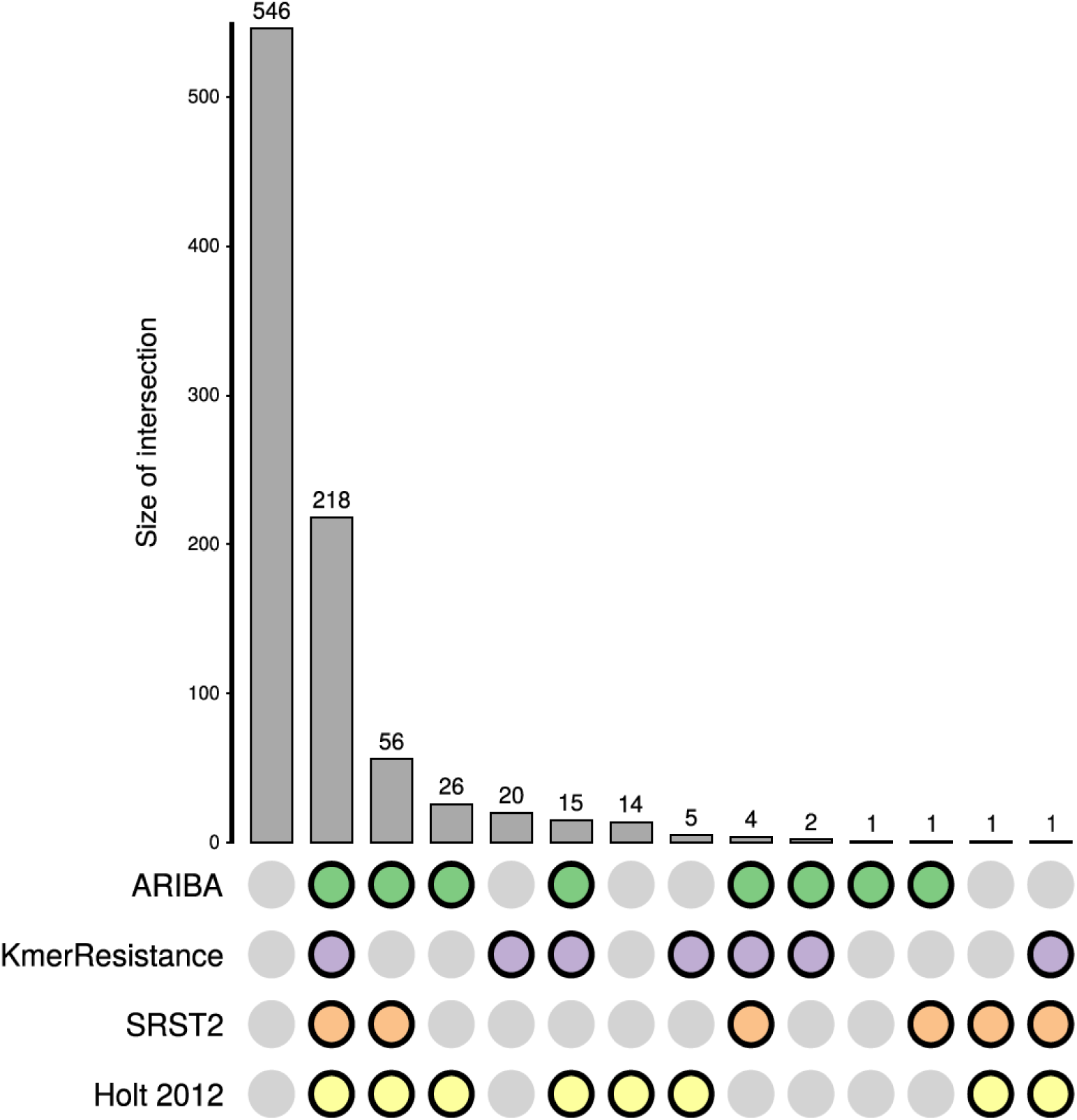
Concordance between AMR calling methods on the S. *sonnei* data. A coloured dot indicates which methods were in agreement. The first column illustrates where no resistance mechanisms were predicted.

For the 146 calls where there were discrepancies between the methods, we observed some general trends explaining most of the discordance. First, neither KmerResistance nor SRST2 identify specific SNPs conferring resistance, whereas this is possible with ARIBA and Holt *et al.* SNPs in the *gyrA* gene, which cause resistance to quinolone antimicrobials, were found by both ARIBA and the manual method of Holt *et al.* in 22 isolates. Second, there were 20 cases where a resistance gene was called by KmerResistance, but not by any other method. Upon further investigation, KmerResistance reported these genes at a low coverage (1.4 - 4.9X). Third, although KmerResistance appears to be the best at detecting genes present at very low coverage, it made fewer calls of genes at higher coverage than ARIBA and SRST2 (Supplementary Figure S8). For example, in isolate ERR028689 *dfrA1* is found at 31X coverage by ARIBA and at 37X by SRST2, but is not reported by KmerResistance. When partial matches are allowed by SRST2 and ARIBA, there are no calls made only by KmerResistance (Supplementary Figure S9). However, less stringency could result in falsepositive calls. A full report of the calls made by each method for each antimicrobial and isolate examined is in Supplementary Table S5.

There were only two cases where ARIBA did not match any other method. These involved either differences in identifying SNPs, or a large insertion into an AMR gene. In the first, ARIBA differed from the results reported in [23]for samples ERR028676 and ERR028677 when identifying SNPs in the *gyrA* gene that confer resistance to quinolone drugs. ARIBA was confirmed to be correctly reporting the SNPs in each sample by analysing the mapped reads, as described in Supplementary Material.

The second case relates to streptomycin resistance, one mechanism for which requires the presence of both the *strA* and *strB* genes. Sample ERR024606 has an insertion into the AMR gene *strA*, which renders it non-functional. The *strA* gene was called as present by SRST2 with high confidence and a depth of 150X, and at 179X by KmerResistance, and at 100% coverage by Holt *et al.* However, ARIBA correctly characterised *strA* as not functionally present as it did not assemble into a single contig; this was manually confirmed to be due to the insertion of *dfrA14* into the middle of *strA* (Supplementary Figure S10), similar to that described previously [27]. We found a second instance of an insertion disrupting an AMR gene, in this case *strB* (Supplementary Figure S11) in isolate ERR028673, and again ARIBA made the correct call. We note that KmerResistance also made the correct AMR call for streptomycin for this isolate, but only because although it called *strB*, it did not call *strA* (called at 80X and 101X by ARIBA and SRST2 respectively).

### Neisseria gonorrhoeae

The sexually-transmitted pathogen *N. gonorrhoeae* is under strict public health surveillance because isolates resistant to the first-line antimicrobials, azithromycin (AZM) and the extended spectrum cephalosporins (ESCs; i.e., cefixime and ceftriaxone) have been reported worldwide. Here, we use a combined collection of 1,517 *N. gonorrhoeae* isolates to illustrate some of the extended features of ARIBA, including the creation and use of customised AMR databases, identification of resistance mutations (SNPs and deletions) in coding and non-coding regions and identification of heterozygous resistance mutations in multicopy rRNAs. The data are from five recent studies [28–32] (Supplementary Table S6) that include phenotypic data on resistance to four antimicrobials. We note that this example is intended to be for illustrative purposes only, not an in-depth analysis of gonococcal AMR determinants.

First, we created a custom database of gonococcal AMR determinants (Supplementary Table S7). Unique alleles for each gene from the 2016 World Health Organization gonococcus reference collection [33] and five available *N. meningitidis* complete genomes (H44 - GCA_000191445.1; MC58 - GCA_000008805.1; M01-240149 - GCA_000191465.1; FAM18 - GCA_000009465.1; Z2491 - GCA_000009105.1) were included in the database to allow identification of recombinant genes. For the purposes of this example, we concentrate on AZM resistance and associated mutations in the 23S rRNA and the *mtrR* gene, which encodes a repressor to the mtr (multiple transferable resistance) efflux system. Our database includes two 23S mutations, A2045G (A2059G *E. coli* numbering) and C2597T (C2611T *E. coli* numbering), which are linked to high-level [34] and low-level [35] AZM resistance, respectively. For *mtrR*, both a G45D substitution and interruption of the gene have been linked to increased efflux leading to reduced susceptibility to multiple antimicrobials.

Visualisation of the ARIBA results in Phandango allows patterns of the presence and absence of variants to be viewed against a tree of isolates. ARIBA can create a dendrogram of isolates based on the identified resistance variants, so that when visualised in Phandango, isolates are clustered by shared resistance-determinant profiles. Alternatively, a phylogenetic tree based on SNPs in the core genome of the isolates can be provided to Phandango, as in Supplementary Figure S12, making it possible to visualise interactively the distribution of resistance mechanisms across the pathogen population. Based on the occurrences of 23S and *mtrR* variants on independent branches within the phylogenetic tree, it is clear that the variants in our database have emerged multiple times in the gonococcal population. 23S-mediated resistance, in particular, has often emerged but failed to spread, suggesting it may be associated with a fitness cost.

Next, we explored the distribution of the Minimum Inhibitory Concentration (MIC) for AZM in isolates with all AZM-related genetic resistance determinants as identified from our database (Supplementary Figure S13) using the “micplot” function of ARIBA. This function outputs publication-quality images along with pairwise Mann-Whitney U Test p-values and effect sizes. Although, as expected, the 23S mutations in our database show clear evidence of association with resistance, the results for the *mtrR* variants are less clear-cut, being found in both resistant and sensitive isolates. Visualising MICs of combinations of resistance determinants allows improved understanding of causal versus linked AMR determinants, and of combinations of determinants which may produce a cumulative effect. By default, ARIBA micplot draws all observed combinations of variants output by ARIBA against user-provided MIC data, so that linked and combinatorial determinants are easier to identify. Figure 4 shows that when separated from linked 23S mutations, the 45D substitution or interruption of *mtrR* alone showed no increase in MIC relative to isolates without a proposed resistance determinant, consistent with other studies [28].

**Figure 4.**
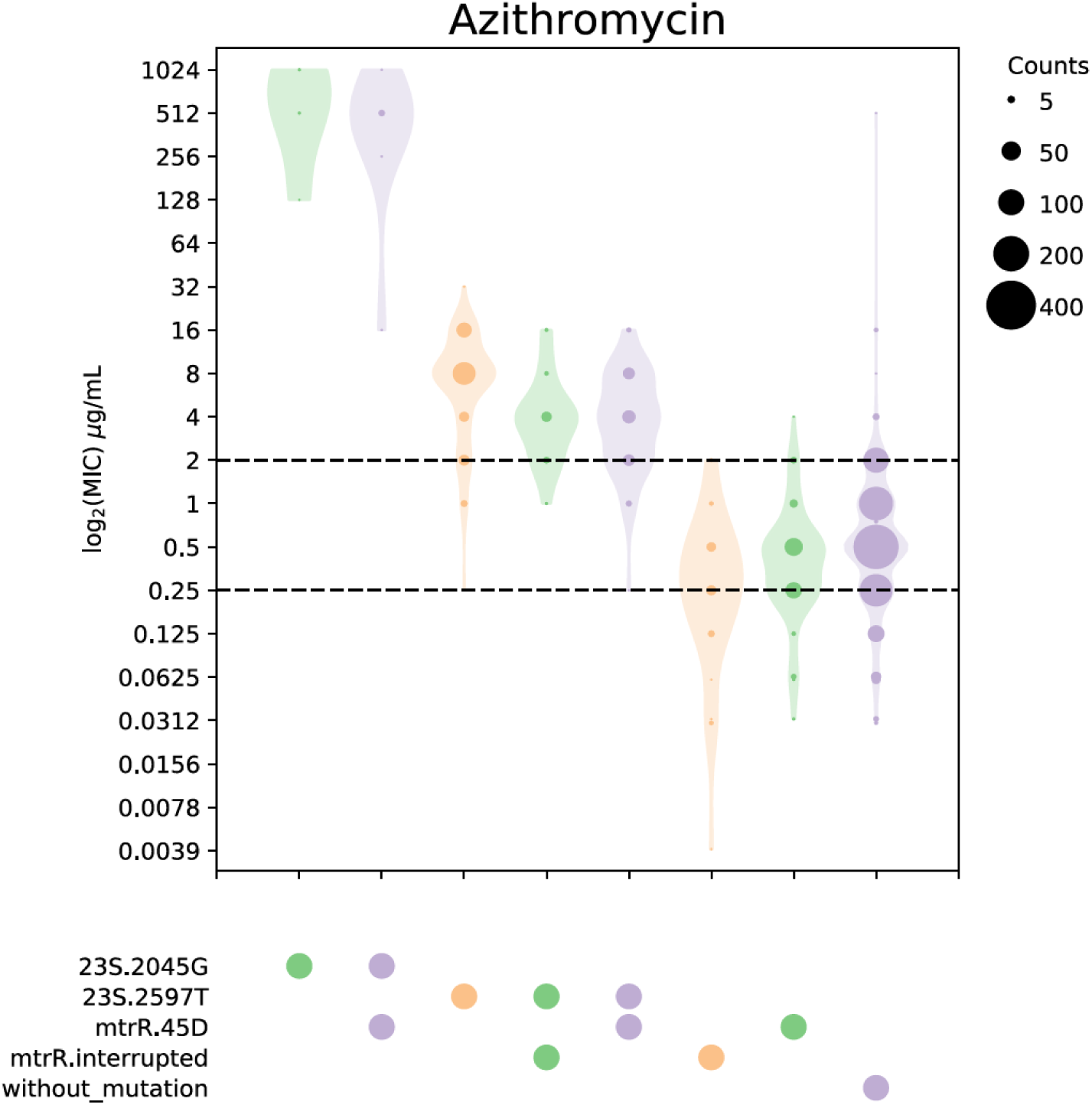
Distribution of MICs (represented on a logarithmic scale) for AZM for all observed combinations of relevant AMR determinants in our custom database. Dotted horizontal lines mark clinical breakpoints. The lower line marks the lowest EUCAST (http://www.eucast.org/clinical_breakpoints/) breakpoint (0.25 μg/mL) and the upper line the post-2005 breakpoint used in the US (2 μg/mL) [28].

Although most of the isolates with the 23S mutations exhibited MICs above the 2μg/mL breakpoint, some would be identified as susceptible if this breakpoint was strictly applied. *N. gonorrhoeae* usually carries four copies of the ribosomal operon. The C2597T mutation can occur in any number of the 23S copies, with increasing number of copies of the mutated allele having been previously associated with increasing MIC [31,36]. ARIBA allows the detection of such heterozygous mutations, which can be important for understanding genotype-phenotype relationships. Supplementary Figure S14 shows how excluding isolates for which the 23S mutations are heterozygous alters the plots in Figure 4, reducing the number of isolates falling below the 2μg/mL breakpoint. Figure 5 shows the percentage of reads (a proxy for the number of gene copies) carrying the mutation, as reported by ARIBA, and its correlation with AZM MIC, confirming that increasing copies of the mutation are linked to increased phenotypic resistance in this dataset.

**Figure 5.**
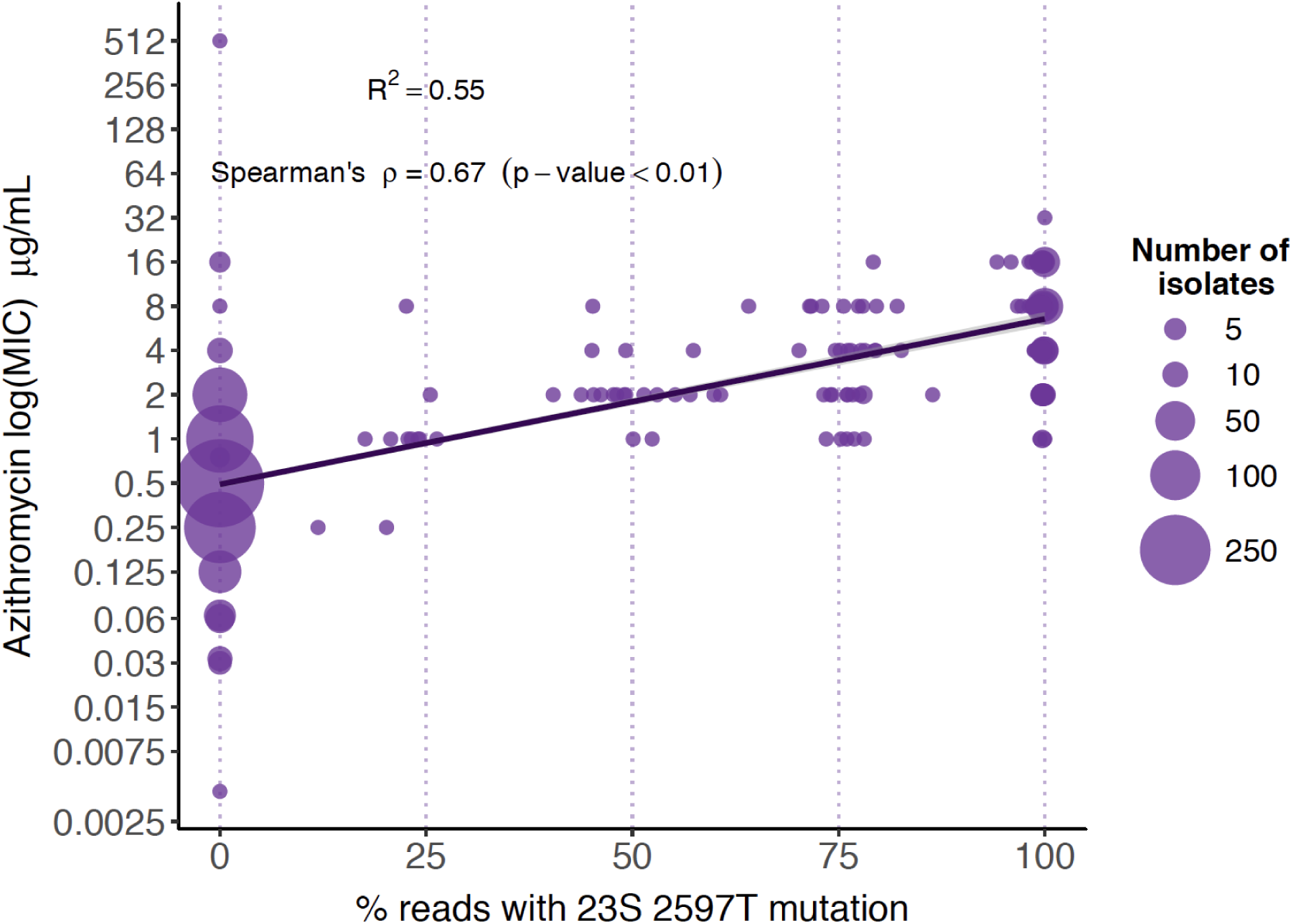
Correlation between the number of alleles containing the 23S C2597T mutation (C2611T in *E. coli*) in AZM-resistant isolates and their MIC values for this antimicrobial.

## Discussion

Increasing antimicrobial resistance threatens to produce untreatable infections, with catastrophic consequences for public health. While new antimicrobials must be developed, we also need to use our current antimicrobials effectively, using those that are appropriate for the resistances and sensitivities of the infection to be treated. One approach to this will be to use rapid genomics-based approaches to predict resistance, and this in turn will rely on fast, accurate and automatable software tools. Here we have developed and implemented a new tool, ARIBA, that not only outperforms existing tools at identifying AMR genes, but also identifies and classifies variants involved in AMR. In addition to supporting common AMR databases, ARIBA was developed to be easily applied with any input reference data. This means that it could be used to identify any sequences of interest, not just those involved in AMR. The use of local assemblies means that ARIBA can determine effectively whether or not an isolate possesses a copy of a gene that is functional or non-functional, unlike other tools, which do not perform this depth of analysis. Further, as showcased on the *N. gonorrhoeae* data, ARIBA reports the presence of variants, interprets their consequences, and identifies the presence of a variant that is known to cause AMR. However, our method is only as good as the quality of the input reference database, and these databases will need to be independently validated, especially if they are intended for clinical use.

## Conclusions

We have developed a new tool, ARIBA, that identifies AMR determinants directly from paired sequencing reads, and have demonstrated a number of ways in which it improves upon existing tools: 1) verifies completeness of acquired resistance genes; 2) identifies known causative resistance SNPs; 3) allows exploration of the association of AMR determinants with user-provided MIC data; 4) identifies SNP frequency in multicopy genes, which has been traditionally difficult to resolve due to the complexities of *de novo* assembly; and 5) generally requires less time and computational resources. Thus, the novel approach of mapping followed by targeted assembly of each reference sequence is fast, efficient and accurate when compared to current methods. Moreover, ARIBA reports significantly more details than existing tools, particularly variant calls, enabling a deeper understanding of the resistance associated with each isolate.

## Methods

### ARIBA

#### Obtaining input data

ARIBA requires reference sequences and, optionally, information about known SNPs. ARIBA supports several public resources, allowing the user to download the data easily and convert it into a form for use with the pipeline. ARG-ANNOT, CARD, PlasmidFinder, ResFinder, VFDB, and the SRST2 version of ARG-ANNOT are currently available. These can be obtained by running the command

~~~
ariba getref name_of_resource output_directory
~~~

Alternatively, input data can be provided by the user. Reference sequences can be coding or non-coding. Coding sequences are subjected to extra checks for consistency, as described in the next paragraph, and extra analysis is performed on them, such as determining if SNPs are synonymous or nonsynonymous. Further, each reference sequence is classified as “presence/absence” where the existence of a sequence within a sample confers antimicrobial resistance, or “variant only” where a known SNP is required for antimicrobial resistance.

#### Preparing input data

All reference data are checked for consistency. First, any coding sequences are required to begin with a start codon, be a complete open reading frame, and end with a stop codon. All reading frames are checked on both strands and any sequence that fails any of the requirements is removed. Any SNP that does not match the reference sequence is also removed. The remaining sequences are clustered using the `cd-hit-est` program from CD-HIT. Since any sequence can be coding or non-coding, and presence/absence or variant only, the sequences generate four disjoint sets of sequences. These sets of sequences are kept separate, with each one clustered individually. The input data is prepared by running the command

~~~
ariba prepareref -f sequences.fasta -m metadata.tsv preparef.out
~~~

where the reference sequences are in the FASTA file `sequences.fasta` and extra information, such as variants, is in the tab-delimited file `metadata.tsv`. These input files are generated automatically when running `ariba getref`.

#### Cluster analysis

Once the reference data are prepared, the main ARIBA pipeline can be run using paired Illumina reads and the reference data as input. The command is

~~~
ariba run prepareref.out reads_1.fq reads_2.fq run.out
~~~

where the directory `prepareref.out` was made when `ariba prepareref` was run. First, the reads are mapped to all of the input sequences (that passed quality filters), using minimap with a *k*-mer length, *k*, of 15 and minimizer window size of 10. A read is considered to be mapped by minimap if: 1) the match length is at least 50 or half of the read length, whichever is smaller; 2) the start position of the match is within 1.1*k* of the start of the read or the reference sequence; 3) the same as for 2), but for the end position of the match. The result is that reads that match completely to the centre of the reference sequence, and reads that overhang the ends of the reference sequence are counted as mapped. The situation is illustrated in Supplementary Figure S1 and explained in detail in the Supplementary Material.

Any read that maps, or whose mate maps, to a reference sequence is allocated to the cluster to which the reference sequence belongs. Note that the same read can be allocated to more than one cluster, for example if two reference sequences lie next to each other in the genome. Each resulting cluster has a set of reference sequences, as determined by CD-HIT, and a set of paired reads.

Each cluster is processed independently as follows (see Supplementary Figure S2). To reduce assembly running time, the reads input to the assembler are randomly downsampled to a maximum of 50X coverage. Since the true reference sequence for this cluster is not yet known, the coverage is (over)estimated using the length of the longest reference sequence for the cluster. The reads are assembled using fermi-lite, which is run using the options `-l x -c y, 10000` where `x` takes the values 6, 15 and `y` takes the values 4, 17, 30, resulting in six distinct assemblies.

The assemblies are compared against all reference sequences from the cluster using nucmer. The best within-cluster nucmer match is identified by maximising for the percent of the reference sequence that is assembled. Ties are broken by taking the highest percent identity, the largest value of `-l` from minimap, and finally the largest value of `-c` from minimap. Next, the assembly contig subsequence from the best nucmer match is compared against all reference sequences (across all clusters). The best match is chosen using the same criteria as for the within-cluster best match, and the corresponding reference sequence is chosen to be the closest reference sequence for this cluster. If the closest reference sequence does not belong to the cluster, then no further analysis is performed and the cluster is not counted as present.

Next, the assembly is compared to the closest reference sequence using the MUMmer suite of programs. The contigs are aligned to the reference sequence using `nucmer`, then SNPS and indels are identified between the sequences using `show-snps`. This information is used to determine the overall success of the assembly, encoded into a bitwise flag (i.e., a single integer). For example, the reference sequence could have a complete match to a single contig. In the case where the reference sequence is a gene, the matching position in the contig is checked for any nonsense mutations. A complete explanation of the flag and the various scenarios it encodes is given in the Supplementary Material. The meaning of a flag `N` can be determined using the command

~~~
   ariba flag N
~~~

which will report a breakdown of the flag `N`.

All reads from the cluster are mapped to the contigs using Bowtie2 and the read depth at each contig position and SNPs are identified using samtools mpileup. Finally, the alignment and variant information is used to generate a summary for this sample, which includes the success of the assembly, whether or not the sample has SNPs of interest and the read depth at those SNPs.

The output of `ariba run` includes a report file containing the summary information of each cluster, plus FASTA files of the assemblies and detailed logging information.

#### Summarizing results

The results of multiple runs of ARIBA across different samples can be summarized by running

~~~
ariba summary out report.*.tsv
~~~

where `report.*.tsv` is a list of reports (each made with a call to `ariba run`). This command generates input files to Phandango, and a CSV file that can be easily viewed in spreadsheet applications. A key output for each sample and cluster is an interpretation of the flag, where how well the matches the reference sequence is summarised as one of: no, partial, fragmented, interrupted, yes_nonunique, or yes (Supplementary Figure S4).

Since Phandango requires a tree, ARIBA determines a rough tree using the contents of its CSV file, which means that it is generated from the calls involving the reference genes and SNPs of interest. The distance between two samples is defined as the number of columns in the CSV file that agree, and an UPGMA tree is generated from the distance matrix using DendroPy [37]. Users may wish to provide their own tree, calculated using sequence-based methods.

## Benchmarking

ARIBA version 2.8.1 was used, together with dependencies Bowtie2 2.2.29, CD-HIT 4.6, MUMmer 3.1 and Python packages dendropy 4.2.0, pyfastaq 3.15.0 [38], pymummer 0.10.2 [39]and pysam 0.10.0 [40]. We used KmerResistance checkout 041bc89b832cf6a3b7629d76b4dffb4c7428caab, and SRST2 0.2.0 with the recommended versions Bowtie2 2.1.0 and samtools 0.1.18. All software was run with the default settings on the Cloud Infrastructure for Microbial Bioinformatics [41]. The complete terminal commands used are in the Supplementary Material.

The details of the 41 *E. faecium* samples with provided accession numbers used and results of gene identification are shown in Supplementary Table S1. In order to sample the *E. faecium* reads at a range of depths, the reads were first mapped to the reference genome CP006620 using bowtie2 version 2.2.29 with the option ‐‐`fast-local`. The depth for each sample was estimated across the *vanB* gene CP006620.1476 by running `samtools depth` with the options `-a -r CP006620:774918-775946` and calculating the resulting mean depth. This was used as an estimate for read depth and the reads were randomly sampled accordingly (this is implemented in the supplementary script `make_read_subsets.pl`) using `fastaq to_random_subset` with a different random seed for each run, producing independent read subsets.

The ARG-ANNOT sequences included with SRST2 were used as reference sequences for the *E. faecium* benchmarking. However, the VanS-B gene, called “47_VanS-B_Gly_VanS-B_1672 no;yes;VanS-B;Gly;AY655721;731-2073;1343” by SRST2, originally from ARG-ANNOT, was missing its final nucleotide A. This was confirmed by comparing with the GenBank record AY655721. It would cause ARIBA to exclude this sequence because the translation into amino acids results in a sequence that does not end with a stop codon. Therefore an “A” was manually added to the end of the sequence before running ARIBA.

The details of the 130 S. *sonnei* samples and AMR calls made by each method are shown in Supplementary Tables S2 and S6. In order to interpret the output of each tool as an AMR call, the following rules were used, where all relevant genes are listed in Supplementary Table S8. A gene was counted as present by ARIBA if ariba summary reported yes or yes_nonunique, present by KmerResistance if it appeared in its output file, and present by SRST2 if it was reported without a “?”.

The focus for the genes of interest for each AMR call were those originally identified and reported in Holt *et al* [23]. Given that the discovery and classification of AMR gene variants is an ongoing process, an AMR gene was called as present if it was either the originally identified gene in Holt *et al*, or in the same CD-HIT cluster. Genes conferring resistance to antimicrobials not examined in the original paper were excluded, as were genes conferring resistance to the antimicrobials examined in the paper but falling in different CD-HIT clusters from the originally identified genes. For each antimicrobial examined, an AMR call for a resistant genotype was identified using the following rules. Ampicillin (Amp): the presence of any gene from a set of *blaTEM*, blaCTX-M and blaOXA genes. Chloramphenicol (Cmp): the presence of any gene from a set of *cat* genes. Nalidixic acid (Nal): the *gyrA* gene present, together with one of the SNPs S83L, D87G, or D87Y. Streptomycin (Str): both of the *strA* and *strB* genes, or one of the *aadA* genes. Sulfonamides (Sul): any gene from the set of *sul1* and *sul2* genes. Tetracycline (Tet): both of *tetA* + *tetR*, or all of *tetA*,*C*,*D*,*R*, where each of the two sets of *tetA* and *tetR* genes are disjoint. Trimethoprim (Tmp): any one of a set of *dfrA* or *dhfr* genes.

The input files and commands run to create the *N. gonorrhoeae* ARIBA resistance database can be found in the supplementary material. Briefly, for each gene all unique alleles from the reference set were saved in multifasta files. For variant-based resistances, alignments were created in Seaview [42,43]by translating to amino acid sequences, aligning with Clustal [42] using default parameters and back translating to nucleotides. For each alignment the `aln2meta` function of ARIBA was used to produce the files required as input to `prepareref`. These were combined, along with the sequence files for presence/absence resistance gene files and `prepareref` run to create the ARIBA database.

To create a phylogenetic tree of all isolates, sequencing reads were aligned to the chromosome of *N. gonorrhoeae* FA1090 (accession number NC_002946) using BWA MEM (version 0.7.12-r1039) [44] with the options to output alignments for unpaired reads and to mark shorter split hits as secondary. Optical duplicates were removed and indels realigned using GATK [45] MarkDuplicates (version 1.127) and indelRealigner (version 3.4-46) respectively, under their default settings. Variant sites were identified from each isolate using samtools (version 1.2) [21] mpileup with options to report DP and DP4 statistics, count orphans, adjust the mapping quality to 50 and increase the maximum depth to 1000, including for indel calling, followed by bcftools (version 1.2) call using a prior of 0.001, a ploidy of 1 and with the option to keep all alternate alleles at variant sites. All sites were further filtered as described previously [46] to produce a multiple sequence alignment. Repeats and prophages in the FA1090 genome were masked from the alignment before variable sites were identified with snp_sites [47]and a neighbour joining phylogenetic tree created with RapidNJ [48]. Interactive visualisation of the phylogenetic tree and ARIBA summary data was carried out in Phandango v0.8.5 [12].

## ARIBA software

ARIBA is open source and available for Linux at https://github.com/sanger-pathogens/ariba under the GNU GPLv3 licence. The implementation is in Python and C++, with low memory usage and short run times compared to other tools (Supplementary Table S2, Supplementary Figure S15).

## Supplementary Files

supplementary.pdf - text and Supplementary Figures S1-15 supporting the main text.

supplementary_tables.xlsx - an Excel file that contains the following supplementary tables.

Table S1. *E. faecium* data (including all accession numbers) and results.

Table S2. Run times and memory usage on the *E. faecium* and S. *sonnei* data.

Table S3. S. *sonnei* data (including all accession numbers) and results.

Table S4. Reference genes manually included in S. *sonnei* analysis.

Table S5. Summary of calls made on S. *sonnei* dataset.

Table S6. *N. gonorrhoeae* data (including all accession numbers).

Table S7. List of antimicrobial genetic determinants included in the *N. gonorrhoeae* ARIBA database. The coding or non-coding nature of the different determinants is indicated along with the cause of resistance.

Table S8. Genes used when determining AMR calls on the S. *sonnei* dataset.

## Authors’ contributions

The methods were devised by MH, AEM and SRH and the ARIBA code was written by MH and AJP. *E. faecium* and S. *sonnei* analyses were performed by MH, AEM, SRH and the *N. gonorrhoeae* analysis by LS-B and SRH. The manuscript was written by all authors.

## Acknowledgements

The authors thank Torsten Seemann, Mark Schultz and the Infection Genomics group at the Sanger Institute for testing and feedback during development of ARIBA, and Sascha Steinbiss for packing the software for Debian.

